# The value of non-motor features and genetic variants of Parkinson’s disease for clustering Lewy body diseases

**DOI:** 10.1101/551507

**Authors:** Olaia Lucas-Jiménez, Ibai Diez, Natalia Ojeda, Naroa Ibarretxe-Bilbao, Javier Peña, Beatriz Tijero, Marta Galdós, Ane Murueta-Goyena, Rocío Del Pino, Marian Acera, Juan Carlos Gómez-Esteban, Iñigo Gabilondo

**Affiliations:** Department of Methods and Experimental Psychology, Faculty of Psychology and Education, University of Deusto, Bilbao, Spain; Department of Neurology, Massachusetts General Hospital, Harvard Medical School, Boston, MA 02114; Neurotechnology Laboratory, Tecnalia Health Department, Derio, Spain; Neurodegenerative Diseases Group, Biocruces-Bizkaia Health Research Institute, Barakaldo, Bizkaia, Spain; Ophthalmology Department, Cruces University Hospital, Barakaldo, Bizkaia, Spain

**Author notes:** Corresponding author: Iñigo Gabilondo, Neurodegenerative Diseases Group, Biocruces-Bizkaia Health Research Institute, Plaza de Cruces 12, Barakaldo (Bizkaia), CP 48903, Spain Corresponding author’s, Corresponding author’s.

**Keywords:** Lewy bodies, Parkinson’s disease, cognition, genetic, clustering analysis

## Abstract

**Introduction:** The use of non-motor Parkinson’s disease (PD) features and genetic PD variants for clustering analyses may refine the phenotypic classification of idiopathic Lewy body (LB) diseases.

**Methods:** One-hundred participants [n=7 E46K-SNCA (n=5 symptomatic and n=2 asymptomatic), n=4 PARK2, n=3 LRRK2, n=8 dementia with Lewy bodies (DLB), n=48 idiopathic PD (iPD), n=30 healthy controls (HC)] underwent a comprehensive evaluation of non-motor and motor PD features. Non-motor features were used to perform a hierarchical clustering analysis with patients and HC using a Scikit-learn toolkit.

**Results:** Clustering analysis suggested three clusters of subjects including Cluster 1 or “*Normal-to-mild*”: young iPD (< 60 years) classified together with most HC and the variable LB burden genetic PD variants (PARK2 and LRRK2) characterized by having normal-to-mild cognitive disabilities and mild-to-moderate motor disability with few axial symptoms; Cluster 2 or “*Mild-to-moderate*”: old iPD patients (>60 years) classified together with the lowest symptomatic E46K-SNCA, PARK2 carriers and HCs, characterizing by having mild-to-moderate cognitive and motor disabilities with few axial symptoms; and Cluster 3 or “*Severe*”: old iPD (>60 years) classified together with all DLB and the most symptomatic E46K-SNCA carriers, characterized by having severe pattern-specific cognitive disabilities (visual attention, perception, processing speed, memory and executive functions) and severe motor PD manifestations with marked axial symptoms.

**Conclusions:** Our study supports the potential value of incorporating genetic PD variants in data-driven iPD classification algorithms and the usefulness of non-motor PD features, especially visual cognition abnormalities, to facilitate the identification of aggressive LB diseases.

## 1. INTRODUCTION

Parkinson’s disease (PD) is a heterogeneous condition with marked variability in terms of clinical presentation and disease progression. Current PD classification perspectives extend far beyond the classically accepted phenotypes based on the predominance, severity and progression of motor features [1,2,3,4]. In fact, non-motor PD clinical features such as cognitive abnormalities, apathy, depression, anxiety, psychotic manifestations, REM sleep behavior disorder, olfactory dysfunction or dysautonomia are now becoming essential to define PD phenotypes, since they are key predictors of disease progression and quality of life [5]. However, the use of exclusively clinical data to define PD subtypes may be insufficient [6]. One of the main reasons for the clinical heterogeneity of PD may be the existence of biologically distinct subtypes of PD. The identification of such biological PD subtypes is of foremost importance since it may favor the development of targeted specific disease-modifying and symptomatic treatments for PD. Therefore, the use of relevant biological data such as genotype information on PD classification paradigms might be extremely useful to refine PD clustering classification algorithms.

Only two publications, both based on data from Parkinson’s Progression Markers Initiative (PPMI), have evaluated the presence of specific genetic PD variants in parkinsonian patients classified through cluster analysis [7,8]. Both studies analyzed the relative distribution of mutation carriers for glucocerebrosidase (GBA) and Leucine-Rich Repeat Kinase 2 (LRRK2) genes among identified PD clusters. In addition, one of such studies [7] applied genetic information as an extra clustering feature using a single genetic indicator termed ‘genetic risk score’ that had been previously calculated for PPMI cohort [9] and which consisted in the summation of the number of 28 common risk variants for PD across 24 loci from genome-wide analyses [10] plus two additional GBA and LRRK2 risk variants detected within PPMI. The key pathological feature of the most aggressive idiopathic PD (iPD) phenotypes is the presence of high burdens of Lewy bodies (LB). However, GBA and specially LRRK2 gene mutation carries have varying degrees of LB deposition in the CNS. Thus, the incorporation to cluster analyses of patients with PD mutations unambiguously associated to severe (E46K-SNCA) [11,12] or absent LB pathology (PARK2) may favor the identification of clinical features and phenotypes linked to severe LB pathology in iPD.

Moreover, the evolution of cognitive impairment in patients with iPD appears to be heterogeneous, and it is important to determine whether specific domain impairment phenotypes can be identified that can characterize specific subgroups of patients which are more sensitive to faster conversion to dementia. Therefore, the objective of this study was to classify idiopathic LB diseases using an innovative cluster analysis with a comprehensive set of non-motor features, including an extensive cognitive evaluation, and involving three genetic PD variants with different degrees of LB pathology.

## 2. METHODS

### 2.1. Study sample and general procedures

One hundred participants were involved in the study: 7 E46K-SNCA, 4 PARK2 and 3 LRRK2 carriers, 8 patients with probable DLB, 48 patients with idiopathic PD (iPD) and 30 healthy controls (HC). From the E46K-SNCA carriers, n=5 were symptomatic with motor and/or non-motor PD manifestations and n=2 were asymptomatic. All HC were recruited to approximately match older symptomatic E46K-SNCA carriers in age and sex. Participants were recruited in the Department of Neurology of Cruces University Hospital and in the PD Biscay Association (ASPARBI). iPD patients fulfilled Parkinson’s UK Brain Bank criteria for the diagnosis of PD and DLB patients the diagnosis of probable DLB by revised criteria for the clinical diagnosis of DLB. There was no family history of parkinsonism in first order relatives of HC, iPD or DLB patients. We did not include participants with progressive neurological disorders other than PD or any medically unstable condition or limiting psychiatric disease. An ophthalmologist excluded participants with any eye condition influencing visual or neuropsychological evaluations. All patients were studied in on-medication and in their optimal on-motor situation to comply with study evaluations. The study protocol was approved by the Basque Clinical Research Ethics Committee. In accordance with the Declaration of Helsinki, all subjects were volunteers and submitted written informed consent prior study participation.

### 2.2. Non-motor symptoms evaluation

#### 2.2.1. Cognitive features

General cognition screening was assessed with Montreal Cognitive Assessment (MoCA)[13]. A comprehensive neuropsychological evaluation of five cognitive domains with the tests recommended by Movement Disorders Society criteria for diagnosis of mild cognitive impairment in PD[14] was also performed: *attention and working memory* with Digit Span Backwards test (DSb)[15] and Trail Making Test A (TMTA)[16]; *executive functions* with Modified Wisconsin Card Sorting Test (Categories) (MWCSTc)[17] and TMTB[16]; *language* with Calibrated Ideational Fluency Assessment [semantic fluency (SF) and letter fluency (LF)][18]; *memory* with Hopkins Verbal Learning Test (HVLT)[19] and Brief Visual Memory Test (BVMT)[20]; *visuospatial functions* with Benton’s Judgment of Line Orientation Test (H-form) (BJLOTH)[21] and Clock Drawing Test order (CDTo)[22]. In addition, we measured *processing speed* with Symbol Digit Modalities Test (SDMT)[23] and Salthouse Perceptual Comparison Test (SPCT)[24]. Outcome variables were converted to z scores to generate composites for each cognitive domain. All composite cognitive domains maintained satisfactory internal consistency with Cronbach α above 0.85 in all cases.

#### 2.2.2. Autonomic, olfactory, ophthalmological and clinical features

We recorded orthostatic hypotension (OHT) with tilt table test, blood pressure recovery time (PRT) following termination of Valsalva maneuver back to baseline (seconds) [25], heart rate response (variability) to deep breathing (HRVdb) (measured as the mean heart rate range in 6 respiration cycles) [26] and olfaction with Brief Smell Identification Test (BSIT) [27]. Binocular low contrast visual acuity (LCVA) [2.5% Sloan charts at 4 meters (Precision Vision, La Salle, IL)] and photopic contrast sensitivity (PCS) [Pelli-Robson chart (Metropia Ltd., Cambridge, UK) at 1 meter with 280 lux chart luminance)] was measured. Moreover, depressive, apathy and fatigue symptoms were evaluated with Geriatric Depression Scale (GDS)[28], Lille Apathy Rating Scale (LARS)[29], and Fatigue Severity Scale (FSS)[30], respectively.

### 2.3. Motor symptoms and PD-related features evaluation

We measured disease duration, Hoehn & Yahr Scale, Unified Parkinson Disease Rating Scale (UPDRS) and levodopa equivalent daily dose (LEDD) [31].

### 2.4. Selection of variables and clustering analysis

To optimize the performance of clustering analyses, we selected the clinical variables that best differentiated patients and HC using Random Forest Classifier (RFC). According to RFC, we chose the following variables for hierarchical clustering analysis: age (demographics); BSIT (olfaction); OHT, PRT, HRVdb (autonomic testing); LCVA and OCS (visual function); GDS (depressive symptoms); DSb and TMTA (attention and working memory); MWCSTc and TMTB (executive functions); SF and LF (language); HVLT and BVMT (memory); BJLOTH and CDTo (visuospatial functions); SDMT and SPCT (processing speed). The former variables were converted to z scores to conduct hierarchical clustering analysis, which was performed including all study subjects, both patients and HC. Features exclusively related to the disease were not included in the analysis and were used post-hoc to compare patients within clusters.

The hierarchical clustering analysis is based on a bottom up approach. Complete linkage criterion was used minimizing the maximum distance between observations of pairs of clusters. k = 3 was selected to offer a good combination of model fit and parsimony. For each identified cluster, we obtained the average (centroid) z score of each variable included to perform hierarchical clustering analysis. All analyses were performed with Scikit-Learning [32] running under Python version 3.6.5.

### 2.5. Data analysis

Normality of data was tested using the Shapiro-Wilk test. Categorical data were analyzed with the Chi-squared (χ2) test. Significant differences in variables were compared using the Analysis of Variance (ANOVA) test or Kruskal-Wallis test and two-tailed t-tests or U-Mann Whitney test for two-group comparisons. Differences between clusters were analyzed with χ2 and ANOVA Tukey-corrected as post-hoc tests for pairwise comparisons. Statistical analyses were performed, using the statistical package SPSS program (IBM SPSS Statistics 22). In addition, to summarize the obtained clusters in graphical representations with only 2 variables defining each subject, we used two different dimensionality reduction techniques: principal component analysis (PCA) and linear discriminant analysis (LDA).

## 3. RESULTS

### 3.1. General description of the whole sample

The general characteristics of study participants are displayed in Table 1. In general terms, they were predominantly male (62.0%) and young (57.4 years). DLB patients were older than the other groups, with statistically significant (*p*<0.05) differences compared to E46K-SNCA carriers, PARK2, iPD and HC. iPD patients were also older than genetic carriers and HC, although differences were statistically significant only when compared to HC. Regarding overall cognitive status (MoCA), both DLB and E46K-SNCA symptomatic carriers were the most affected, and there were not significant differences between them. However, DLB patients showed significant lower cognitive scores compared to E46K-SNCA asymptomatic carriers, iPD patients, PARK2 carriers and HC. In addition, HC showed higher cognitive scores compared to E46K-SNCA symptomatic carriers and iPD patients. In terms of PD-related motor status, most diagnostic categories were comparable with UPDRS III ranging from 23.50 to 27.43 and with Hoehn & Yahr stage between 2 and 2.5, except for E46K-SNCA asymptomatic carriers and DLB (with respectively normal and severely affected motor status).

**Table 1.** Demographical and Parkinson’s disease related data of study participants

|  | **E46K-SNCA symptomatic** | **E46K-SNCA asymptomatic** | **LRRK2** | **PARK2** | **iPD** | **DLB** | **HC** |
| --- | --- | --- | --- | --- | --- | --- | --- |
| No. participants | 5 | 2 | 3 | 4 | 48 | 8 | 30 |
| Age (years) | 52.47<sup>a</sup><br>(12.40) | 44.87<sup>b</sup><br>(15.05) | 53.84<br>(11.17) | 49.91<sup>g</sup><br>(16.39) | 60.37<sup>c,d</sup><br>(8.23) | 73.97<sup>e</sup><br>(7.29) | 51.19<br>(12.68) |
| Education (years) | 15.40<br>(2.70) | 16.00<br>(5.66) | 9.33<br>(3.06) | 11.25<br>(5.12) | 10.33<sup>d</sup><br>(4.15) | 9.75<br>(4.23) | 14.37<br>(4.97) |
| Males, no (%) | 3<br>(60.00) | 1<br>(50.00) | 2<br>(66.70) | 3<br>(75.00) | 30<br>(62.50) | 6<br>(75.00) | 17<br>(56.67) |
| MoCA | 20.80<br>(7.26) | 29.50<sup>b</sup><br>(.71) | 24.00<br>(4.35) | 25.75<sup>g</sup><br>(3.40) | 24.42<sup>c,d</sup><br>(3.07) | 17.62<sup>e</sup><br>(7.52) | 27.67<sup>h</sup><br>(2.39) |
| Disease duration (years) | 7.62<br>(5.19) | --- | 5.10<br>(4.62) | 8.03<br>(10.08) | 6.38<br>(3.96) | 7.98<br>(5.76) | --- |
| Age at disease onset (years) | 44.81<sup>a</sup><br>(10.53) | --- | 48.70<br>(6.59) | 41.85<sup>g</sup><br>(21.79) | 53.95<sup>c</sup><br>(7.88) | 65.94<br>(9.53) | --- |
| UPDRS I | 4.60 | 0 | 1.00<sup>f</sup> | 1.00<sup>g</sup> | 2.28<sup>c</sup> | 5.29 | --- |
|  | (2.07) | (0) | (1.00) | (1.00) | (1.78) | (2.06) |  |
| UPDRS II | 14.60 | 0 | 13.00 | 11.00 | 12.17 | 16.29 |  |
|  | (9.81) | (0) | (10.15) | (6.08) | (6.56) | (5.94) |  |
| UPDRS III | 27.00 | 1.00 | 26.33 | 23.33 | 27.46 | 37.14 |  |
|  | (17.86) | (1.41) | (8.96) | (10.07) | (11.20) | (13.74) |  |
| UPDRS IV | 4.80 | 0 | 6.00 | 3.00 | 4.48 | 2.43 |  |
|  | (4.38) | (0) | (4.58) | (3.46) | (3.71) | (2.57) |  |
| Hoehn & Yahr (no. patients) |  |  |  |  |  |  | --- |
| Stage 0 | 1 | 2 | 0 | 0 | 0 | 0 |  |
| Stage 1 | 0 | 0 | 0 | 1 | 4 | 0 |  |
| Stage 1.5 | 0 | 0 | 0 | 1 | 6 | 0 |  |
| Stage 2 | 1 | 0 | 2 | 1 | 15 | 1 |  |
| Stage 2.5 | 2 | 0 | 1 | 0 | 15 | 3 |  |
| Stage 3 | 1 | 0 | 0 | 1 | 7 | 4 |  |
| Stage 4 | 0 | 0 | 0 | 0 | 1 | 0 |  |
| LEDD (mg/day) | 716.30<br>(635.99) | — | 713.00<br>(357.98) | 380.00<br>(233.88) | 648.23<br>(395.23) | 570.00<br>(347.33) | --- |

Results are presented as means (standard deviations) unless otherwise specified. Statistically significant ( $p < 0.05$ ) results for pairwise group comparisons with $\chi^2$ (proportions) or one-way ANOVA (means) are represented as: a. E46K-SNCA symptomatic vs DLB; b. E46K-SNCA asymptomatic vs DLB; c. iPD vs DLB; d. iPD vs HC; e. DLB vs HC; f. LRRK2 vs DLB; g. PARK2 vs DLB; h. E46K-SNCA symptomatic vs HC. Abbreviations: E46K-SNCA: E46K mutation in alpha-synuclein gene; “E46K-SNCA symptomatic”: symptomatic carriers of E46K mutation in alpha-synuclein gene; “E46K-SNCA asymptomatic”: asymptomatic carriers of E46K-SNCA; LRRK2: carriers of Leucine-Rich Repeat Kinase 2 gene mutations; PARK2: carriers of mutations in Parkin gene; iPD: idiopathic Parkinson’s disease; DLB: dementia with Lewy bodies; HC: healthy controls; MoCA: Montreal Cognitive Assessment; UPDRS: Unified Parkinson Disease Rating Scale; LEDD: Levodopa equivalent daily dose.

### 3.2. General description of clusters

The hierarchical clustering analysis distributed study participants in three clusters: cluster 1 “*normal-to-mild*” (N=45), cluster 2 “*mild-to-moderate*” (N=23) and cluster 3 “*severe*” (N=32). Table 2 shows the distribution of participants in the three clusters according to their diagnostic category and it also presents cluster differences for all variables included in the clustering. Table 3 shows cluster differences in other relevant variables and PD features not used for hierarchical clustering, which in fact were statistically significant for most comparisons, even after excluding HC.

**Table 2.** Clusters of participants and their differences for variables used in HCA

|  |  | **Cluster 1**<br>*Normal-to-mild* | **Cluster 2**<br>*Mild-to-moderate* | **Cluster 3**<br>*Severe* | **F/ Chi** | **p** |
| --- | --- | --- | --- | --- | --- | --- |
| No. participants | All | 45 | 23 | 32 | --- | --- |
|  | HC | 24 | 6 | 0 | --- | --- |
|  | E46K-SNCA asymptomatic | 2 | 0 | 0 | --- | --- |
|  | E46K-SNCA symptomatic | 1 | 1 | 3 | --- | --- |
|  | LRRK2 | 2 | 0 | 1 | --- | --- |
|  | PARK2 | 2 | 1 | 1 | --- | --- |
|  | iPD | 14 | 15 | 19 | --- | --- |
|  | DLB | 0 | 0 | 8 | --- | --- |
| Demographics | Age (years) | 47.83 (10.22) | 61.37 (4.90) | 67.95 (6.77) | .59.60 | .000<br>a,b,c |
| Cognition | Attention & working memory | .67 (.52) | -.12 (.58) | -.87 (.54) | 72.58 | .000<br>a,b,c |
|  | Executive | .59 (.38) | -.06 (.61) | -.76 (.88) | 40.99 | .000<br>a,b,c |
|  | Language | .59 (.48) | -.02 (.74) | -.80 (.66) | 46.93 | .000<br>a,b,c |
|  | Memory | .68 (.49) | .08 (.47) | -1.07 (.56) | 105.23 | .000<br>a,b,c |
|  | Visuospatial | .55 (.33) | .10 (.44) | -.77 (.85) | 48.34 | .000<br>a,b,c |
|  | Processing speed | .73 (.60) | -.01 (.49) | -1.05 (.50) | 91.91 | .000<br>a,b,c |
| Neuropsychiatry | Depression (GDS) | 1.27 (1.54) | 3.04 (2.77) | 4.16 (3.64) | 11.43 | .000<br>b,c |
| Olfaction | BSIT (#correct out of 12) | 9.75 (1.60) | 7.61 (2.53) | 5.43 (2.63) | 33.90 | .000<br>a,b,c |
| Dysautonomia | Valsalva PRT (sec) | 2.17 (1.66) | 3.17 (2.82) | 4.14 (4.87) | 3.40 | .037<sup>b</sup> |
|  | HRVdb | 1.05 (.10) | .98 (.08) | .90 (.05) | 18.78 | .000<br>a,b,c |
|  | Orthostatic Hypotension | 8 (32%) | 5 (23%) | 5 (45%) | 2.85 | .063<sup>d</sup> |
| Vision | LCVA (#correct letters) | 37.97 (5.29) | 31.08 (5.79) | 15.45 (12.65) | 65.65 | .000<br>a,b,c |
|  | PCS (logMar) | 2.10 (.09) | 1.93 (.10) | 1.84 (.14) | 47.73 | .000<br>a,b,c |

Results are presented as means (standard deviations) unless otherwise specified. Pairwise cluster comparisons were performed using with post-hoc $\chi^2$ tests or Tukey's range test, respectively. Statistically significant ( $p < 0.05$ ) results for pairwise cluster comparisons with $\chi^2$ (proportions) or one-way ANOVA (means) are represented as: "a" cluster 3 versus cluster 2; "b" cluster 3 vs cluster 1; "c" cluster 2 vs cluster 1; "d" tendency for statistical significance ( $p = 0.063$ ) for cluster 2 vs cluster 1. Abbreviations: E46K-SNCA: E46K mutation in alpha-synuclein gene; "E46K-SNCA asymptomatic": asymptomatic
carriers of E46K-SNCA; “E46K-SNCA symptomatic”: symptomatic carriers of E46K mutation in alpha-synuclein gene; LRRK2: carriers of Leucine-Rich Repeat Kinase 2 gene mutations; PARK2: carriers of mutations in Parkin gene; iPD: idiopathic Parkinson’s disease; DLB: dementia with Lewy bodies; HC: healthy controls; GDS: Geriatric Depression Scale; BSIT: Brief Smell Identification Test; PRT: blood pressure recovery time; HRVdb heart rate response (variability) to deep breathing; LCVA: binocular low contrast visual acuity (2.5% Sloan charts); PCS: photopic contrast sensitivity (Pelli-Robson chart). See methods for further details on used protocols for clinical evaluations.

**Table 3.**
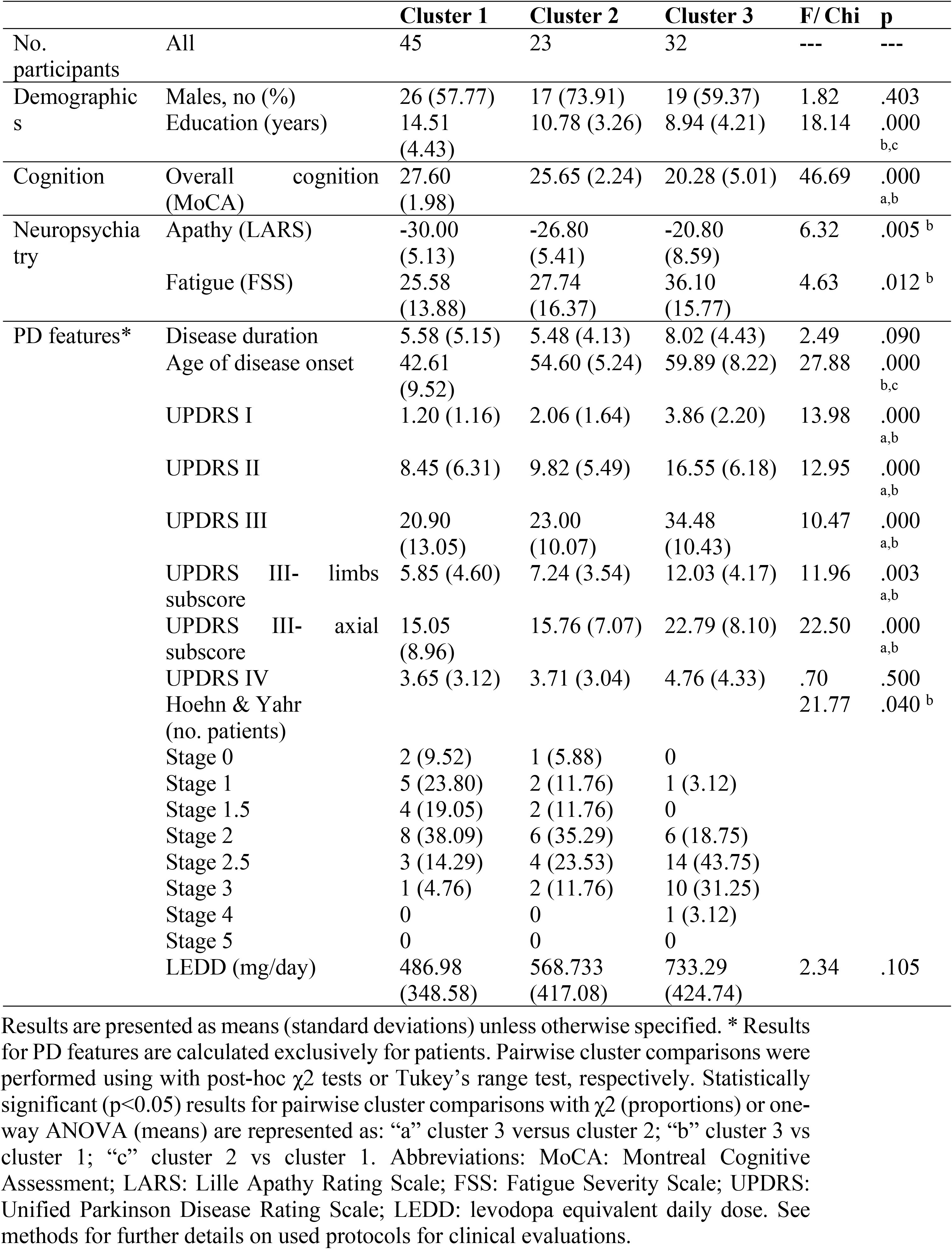
Differences in PD features and variables not included in HCA

Overall, clustering analysis separated participants according to a non-motor manifestations severity grading in which cluster 3 “*severe*” accounted for the most affected subjects, cluster 2 “*mild-to-moderate*” for those in an intermediate clinical status and cluster 1 “*normal-to-mild*” for those that were less disabled. Cluster 3 “*severe*” included old (>60 years) iPD patients (40%) classified together with all DLB and the most symptomatic E46K-SNCA carriers, corresponding to the most aggressive idiopathic and genetic LB diseases (100% of DLB and 60% of E46K-SNCA symptomatic carriers) and it did not include any HC. Cluster 2 “*mild-to-moderate*” included old (>60 years) iPD patients (32%) classified together with one symptomatic E46K-SNCA carrier and with HC (20% and 20% respectively). Finally, Cluster 1 “*Normal-to-mild*” included young (< 60 years) iPD patients (28%) classified together with most HC (80%) and the absent and variant LB burden genetic PD variants (PARK2 and LRRK2).

In Figure 1-A, we can observe the PCA and LDA representation plots with the individual distribution of study participants according to their performance in the non-motor clinical features used for clustering analysis. Accordingly, we may observe comparable PCA and LDA coordinates within cluster 3 for DLB and E46K-SNCA symptomatic carriers and within cluster 1 for PARK2, LRRK2, asymptomatic E46K-SNCA carriers and HC. Interestingly, the E46K-SNCA symptomatic carrier with an isolated pure autonomic failure was situated in the moderate phenotype cluster (cluster 2) and the young E46K-SNCA carrier with mild PD manifestations was situated together with asymptomatic E46K-SNCA carriers in the mild phenotype cluster (cluster 1).

**Fig 1.**
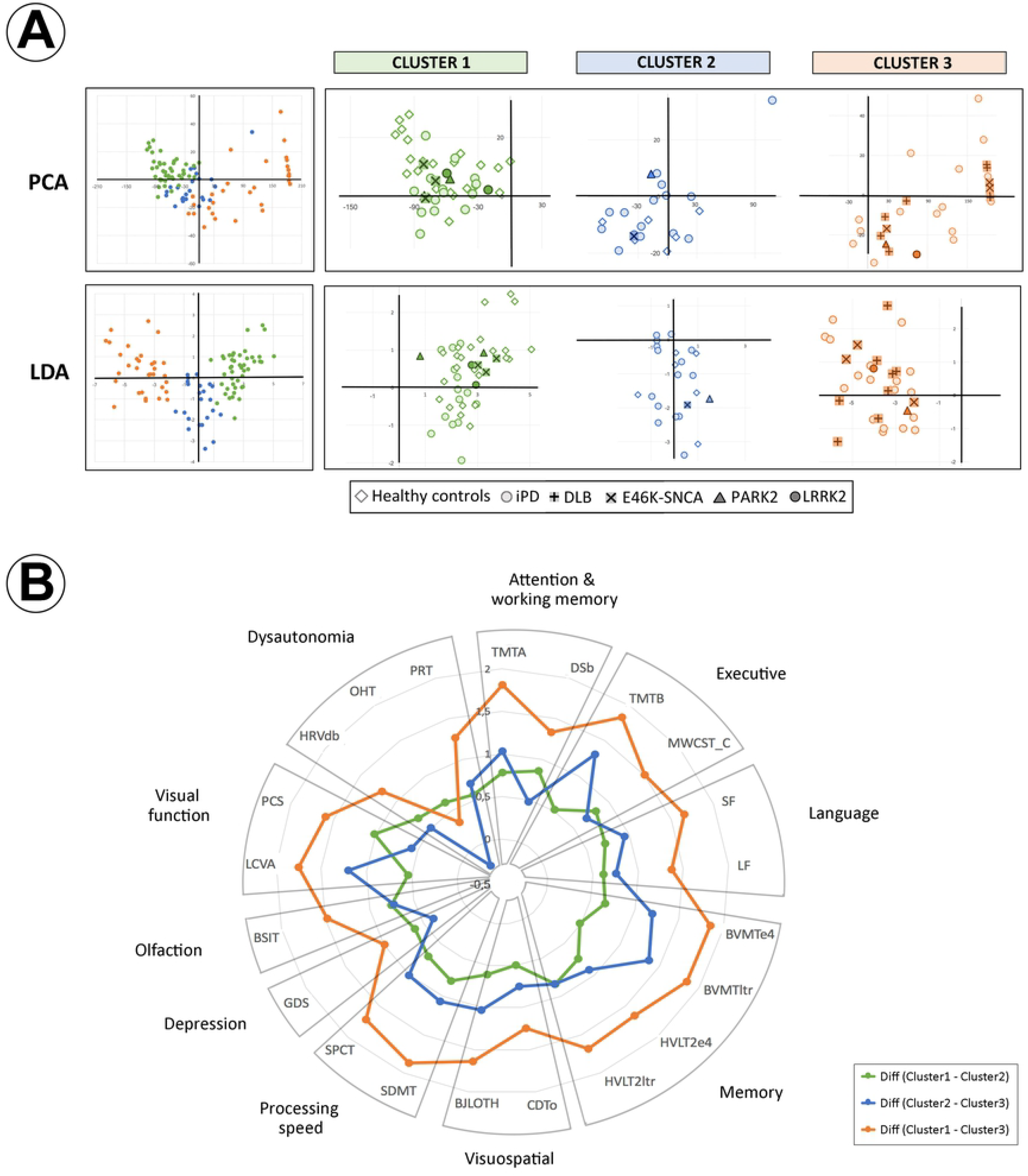
Biplots and radial plot of the three identified clusters in the hierarchical clustering analysis. **a. Biplots for principal component analysis and linear discriminant analysis.** PCA and LDA biplots representing individual distribution of study participants according to their performance in the non-motor clinical features used for HCA. Points that are close together in PCA and LDA biplots correspond to observations that have similar scores on the components displayed in the plot. See main text (Results section) for further interpretation of results in the figure. Abbreviations: PCA: Principal Component Analysis; LDA: Linear Discriminant Analysis; iPD: idiopathic Parkinson’s Disease; DLB: Dementia with Lewy bodies; E46K-SNCA: carriers of E46K mutation of alpha-synuclein gene (SNCA); PARK2: carriers of mutations in Parkin gene; LRRK2: carriers of Leucine-Rich Repeat Kinase 2 gene mutations. **b. Radial plot showing differences in clinical features between identified clusters.** The magnitude of pairwise cluster differences in average z scores (centroids) for each clinical variable included to perform HCA are displayed in a radial plot. See main text (Results section) for further interpretation of results in the figure. Abbreviations: TMTA: Trail Making Test A; DSb: Digit Span Backwards test; MWCST_C: Modified Wisconsin Card Sorting Test (Categories); TMTB: Trail Making Test B; SF: Calibrated Ideational Fluency Assessment semantic fluency; LF: Calibrated Ideational Fluency Assessment letter fluency; HVLT: Hopkins Verbal Learning Test; BVMT: Brief Visual Memory Test; BJLOTH: Benton’s Judgment of Line Orientation Test (H-form); CDTo: Clock Drawing Test order; SDMT: Symbol Digit Modalities Test; SPCT: Salthouse Perceptual Comparison Test; GDS: Geriatric Depression Scale; BSIT: Brief Smell Identification Test; LCVA: low contrast visual acuity; PCS: Photopic Contrast Sensitivity; HRVdb: heart rate response (variability) to deep breathing; OHT: Orthostatic Hypotension; PRT: blood Pressure Recovery Time.

#### 3.2.1. Non-motor symptoms

Regarding demographical variables, patients in cluster 1 were markedly younger, with higher education and earlier disease onset as compared to those in other clusters. In terms of non-motor features, we also observed significant differences for all pairwise cluster comparisons in neuropsychological, olfactory, autonomic and visual features, with a gradient in which cluster 1 was the least affected, cluster 2 the clinically intermediate one and cluster 3 the most severely affected. Regarding neuropsychiatric manifestations, patients in cluster 1 were significantly less depressed, apathetic and fatigued as compared to cluster 1 and cluster 2, who had no significant differences. To identify specific patterns of non-motor features in the clusters, we plotted the magnitude of pairwise cluster differences in average z scores (centroids) for each clinical variable included to perform hierarchical clustering analysis (Fig. 1-B). Interestingly, when cluster 3 was compared with cluster 1 and with cluster 2, we observed a peculiar pattern revealing marked differences in neuropsychological tests (visual attention, perception, processing speed and memory and executive functions), that was not present when differences between cluster 1 and cluster 2 were evaluated.

#### 3.2.2. Motor symptoms and PD-related features

Regarding PD-related features, patients in cluster 1 presented earlier disease onset as compared to those in other clusters. Although patients in cluster 3 had longer disease duration as compared to participants in other clusters, the cluster differences for this variable were not statistically significant. Thus, we did not include disease duration as a covariate in subsequent analyses. The severity of motor PD manifestations was markedly higher in patients from cluster 3 as compared to those from clusters 1 and 2 (*p*<0.05), while it was comparable between cluster 1 and cluster 2. Although there was a progressive increase of LEDD as motor manifestations increased from cluster 1 to cluster 3, cluster differences in LEDD were not statistically significant. Interestingly, axial motor manifestations were significantly more severe for patients in cluster 3 as compared to cluster 1 and cluster 2.

## 3. DISCUSSION

To the best of our knowledge, this is the first attempt to explore the heterogeneity of idiopathic LB diseases using a data driven classification approach with an extensive set of cognitive and other non-motor features. Additionally, we included three pathophysiological and clinically distinct genetic PD variants known for having different degrees of LB disease in the CNS: PARK2 (absent LB pathology), LRRK2 (variable LB pathology) and E46K-SNCA (severe LB pathology). Our clustering analyses identified three groups of patients classified irrespective of disease duration: Cluster 1 or “*Normal-to-mild*” included young iPD, most HC and the lowest LB burden genetic PD variants (PARK2 and LRRK2) characterized by having normal-to-mild cognitive disabilities and mild-to-moderate motor disability with few axial symptoms; Cluster 2 or “*Mild-to-moderate*” included old iPD patients, one symptomatic E46K-SNCA carrier and HC, characterizing by having mild-to-moderate cognitive and motor disabilities with few axial symptoms; Cluster 3 or “*Severe*” included old iPD, all DLB and the most symptomatic E46K-SNCA carriers, characterized by having severe pattern-specific cognitive disabilities (visual attention, perception, processing speed, memory and executive functions) and severe motor PD manifestations with marked axial symptoms. According to our HCA, the clinical pattern observed in cluster 3 may correspond to iPD with higher LB pathology burden since all DLB and most symptomatic E46K-SNCA carriers were included in cluster 3. Similar results were found in other studies that showed a cluster of PD patients with severe motor symptoms, orthostatic hypotension, cognitive impairment, REM sleep behavior disorder, and neuropsychiatric symptoms [4].

Until now, few publications have used clustering analyses to identify PD subtypes based on non-motor symptoms [3,33–35]. In such studies, three or four subtypes of PD patients were usually identified, including the phenotypes “old age onset and rapid disease progression” and “young age onset and slow disease progression”, which may correspond respectively to the PD phenotypes identified in the present work as cluster 3 and cluster 1. Furthermore, our study suggested the existence of a specific PD phenotype (cluster 3) that was identified irrespective of disease duration and which included marked axial motor manifestations and severe non-motor disability.

Moreover, in our study, Valsalva PRT, a recognized early biomarker of dysautonomia [25], was an important variable to differentiate the three clusters. In line with the concept of dysautonomia and aggressive LB disease phenotypes, Kaufmann et al. [36] reported the natural history of 100 patients with pure autonomic failure (a rare synucleinopathy characterized by the presence of severe isolated dysautonomia). In the study, after 4 years of follow-up most of the patients that phenoconverted to a diffuse synucleinopathy did so to DLB, which is characterized by the collection of clinical features mainly represented in cluster 3 of our study. Considering that cluster 3 included not only all DLB patients but also the most symptomatic E46K-SNCA carriers, our findings support the idea that iPD patients with the aforementioned motor and non-motor phenotype may correspond to aggressive diffuse LB diseases. Remarkably, although visuospatial cognitive disability was not the most prominent cognitive feature in cluster 3, most of the non-motor features of cluster 3 are predominantly related to visual cognition. This concept connects with previous evidence supporting that early visual cognitive dysfunction is one of the main predictors for the development of cognitive disability in PD [37, 38]. Processing speed is a cognitive domain that is not usually included in the definition criteria for PD-MCI, but numerous studies have found that the presence of processing speed alterations are widely present in PD, being associated with impairments in daily living activities [39].

However, some limitations should be considered in our study. First, although the inclusion of genetic PD variants in the clustering analysis is one of the highlights of the present work, the sample size of each genetic group is small, which may limit the statistical power and generalizability of study results. However, it is important to remark that SNCA-linked mutations are considered a rare condition as they are limited to specific families and series around the world and their study is a unique opportunity to improve our understanding of the pathophysiology underlying the different phenotypes of LB diseases [11]. Second, it is important to consider the limitations regarding the intrinsic variability of clustering analyses. Third, the mutual relationship between processing speed, executive functions, visuospatial cognition and their decline with chronological age may have influenced the classification of study participants. Fourth, although we performed a comprehensive evaluation of several motor and non-motor clinical features, we were not able to assess the presence of RBD in our study population, an important feature known to be highly predictive of the development of aggressive LB diseases. Moreover, since statistically significant cluster differences persisted even after removing the effect of chronological age on cognitive performance, we cannot completely rule out the effect of age in our analyses, which is in fact a determinant feature conditioning neurodegeneration in PD. Finally, our results are based on cross-sectional data. Therefore, further longitudinal studies are required to investigate whether members of different clusters reveal various course of progression or outcome over time.

In conclusion, our clustering analysis, based on an extensive set of non-motor features and including HC, iPD, DLB and three pathophysiological and clinically distinct genetic PD variants differentiated three clusters of profiles in LB diseases. First, “*Normal-to-mild*” cluster with young iPD patients, most HC and the lowest LB burden genetic PD variants (PARK2 and LR RK2) characterized by having normal-to-mild cognitive disabilities and mild-to-moderate motor disability with few axial symptoms; “*Mild-to-moderate*” cluster with old iPD patients classified together with the lowest symptomatic E46K-SNCA carrier and with HCs, characterizing by having mild-to-moderate cognitive and motor disabilities with few axial symptoms; and “*Severe*” cluster with old iPD classified together with all DLB and the most symptomatic E46K-SNCA carriers, characterized by having severe pattern-specific cognitive disabilities (visual attention, perception, processing speed, memory and executive functions) and severe motor PD manifestations with marked axial symptoms. Hence, our study with genetic PD patients supports the potential value of quantifying non-motor PD features in the clinical setting, particularly visual cognition abnormalities, to help in the identification of those iPD patients at higher risk of developing an aggressive diffuse LB diseases.

## Acknowledgment

We would like to thank ASPARBI and all of the participants in the study for making this research possible.

